# Regulation of brain aging by neutral sphingomyelinase 2

**DOI:** 10.1101/2021.06.08.445892

**Authors:** Zhihui Zhu, Zainuddin Quadri, Simone M. Crivelli, Ahmed Elsherbini, Liping Zhang, Priyanka Tripathi, Haiyan Qin, Emily Roush, Stefka D. Spassieva, Mariana Nikolova-Karakashian, Timothy S. McClintock, Erhard Bieberich

## Abstract

We have shown that deficiency of neutral sphingomyelinase 2 (nSMase2), an enzyme generating the sphingolipid ceramide, improves memory in adult mice. Here, we performed sphingolipid and RNA-seq analyses on the cortex from 10 month-old nSMase2-deficient (*fro/fro*) and heterozygous (+/*fro*) mice. *fro*/*fro* cortex showed reduced levels of ceramide, particularly in astrocytes. Differentially abundant transcripts included several functionally related groups, with decreases in mitochondrial oxidative phosphorylation and astrocyte activation transcripts, while axon guidance and synaptic transmission transcripts were increased, indicating a role of nSMase2 in oxidative stress, astrocyte activation, and cognition. Experimentally induced oxidative stress decreased the level of glutathione (GSH), an endogenous inhibitor of nSMase2, and increased immunolabeling for ceramide in primary +/*fro* astrocytes, but not in *fro/fro* astrocytes. β-galactosidase activity was lower in 5-weeks old *fro/fro* astrocytes, indicating delayed senescence due to nSMase2 deficiency. In *fro/fro* cortex, levels of the senescence markers C3b and p27, and the proinflammatory cytokines interleukin 1β, interleukin 6, and tumor necrosis factor α were reduced, concurrent with 2-fold decreased phosphorylation of their downstream target, protein kinase Stat3. RNA and protein levels of the ionotropic glutamate receptor subunit 2b (Grin2b or NR2B) were increased by 2-fold, an effect known to enhance cognition. This was consistent with 3.5-fold reduced levels of exosomes carrying miR-223-3p, a micro-RNA downregulating Grin2b. In summary, our data show that nSMase2 deficiency prevents oxidative stress-induced elevation of ceramide and secretion of exosomes by astrocytes that suppress neuronal function, indicating a role of nSMase2 in the regulation of neuroinflammation and cognition during brain aging.

**Significance statement:** Oxidative stress is associated with brain aging and cognitive decline. The underlying mechanism how oxidative stress impairs brain function is still not clear. We provide evidence that oxidative stress increases ceramide in astrocytes, which is prevented by deficiency of nSMase2, an enzyme that is activated by oxidative stress and generates ceramide from sphingomyelin. Mass spectrometric and transciptomic (RNA-seq) analyses show that in middle aged (10-month old) mouse cortex, nSMase2 deficiency reduces ceramide and increases expression of genes important for synaptic transmission and cognition. Therefore, our data show that oxidative stress-induced activation of nSMase2 and generation of ceramide is significant for cognitive decline during aging.

## Introduction

The loss of cognition induced by the development of dementia represents one of the main pathological symptoms of human aging. Aging-related cognitive decline in humans can begin as early as middle age (45-65 years old) (Singh-Manoux et al., 2012). Cognitive decline related to the aging brain is associated with oxidative stress and neuroinflammation (Hajjar et al., 2018). Astrocytes, the most versatile cells in the central nervous system, play an important role in brain aging. The oxidative stress-preventing function of astrocytes is mainly due to their ability to generate endogenous anti-oxidants and regulate neuroinflammation (Lee et al., 2010). Most recently, it was shown that in reactive astrocytes, the level of the sphingolipid ceramide was increased (de Wit et al., 2019). Studies from our laboratory and others have shown that astrocytes increase the level of ceramide due to the activation of neutral sphingomyelinase 2 (nSMase2) by amyloid beta (Aβ) peptide and proinflammatory cytokines, the levels of both of which are increased during normal aging and in Alzheimer’s disease (AD) (Singh et al., 1998; Nikolova-Karakashian et al., 2008; Filippov et al., 2012; Wang et al., 2012; Gu et al., 2013; Dinkins et al., 2015; Dinkins et al., 2016; de Wit et al., 2019; Crivelli et al., 2020). We discovered that nSMase2 inhibition or deficiency ameliorated AD pathology in the 5XFAD mouse model, including cognitive improvement (Dinkins et al., 2014; Dinkins et al., 2016). Surprisingly, cognition was also improved in nSMase2-deficient mice without AD (Dinkins et al., 2016). These data suggested that downregulation of ceramide generation is beneficial for cognitive performance and prompted us to investigate the function of nSMase2 and ceramide during normal aging of the brain.

Activation of nSMase2 and in turn, ceramide generation is induced by oxidative stress (Nikolova-Karakashian et al., 2008). Oxidative stress is downregulated by astrocytes that are the main source for glutathione (GSH), an endogenous antioxidant, the level of which is strongly correlated with cognitive performance in the aging brain (Hajjar et al., 2018). The level of GSH determines the activation state of nSMase2 in hepatocytes (Liu and Hannun, 1997; Rutkute et al., 2007). Whether GSH also regulates nSMase2 in astrocytes is not known. If the GSH level drops below 5-8 mM, a concentration found in hepatocytes and astrocytes, nSMase2 is no longer inhibited and generates ceramide (Lee et al., 2010; McBean, 2017). In neurons, activation of nSMase2 by reduction of GSH is not likely since the neuronal GSH concentration (< 1 mM) is far below the range required for nSMase2 inhibition (McBean, 2017). Therefore, inhibition of nSMase2 by GSH is suggested to be critical for the regulation of ceramide levels in astrocytes, but not in neurons. Ceramide levels in nSMase2-deficient (*fro/fro*) astrocytes are expected to be insensitive to a decrease of GSH levels either as a result of aging or experimentally induced by oxidants.

In this study, we analyzed the ceramide and mRNA levels in the cortex of middle-aged (10 month-old) nSMase2-deficient (*fro/fro*) mice and control litermates (+/*fro* and/or wild type). In addition, we determined the effect of experimentally induced oxidative stress *in vitro* and analyzed proinflammatory cytokines and downstream cell signaling pathways activated by ceramide *in vivo*. Our studies show that nSMase2 deficiency decreased ceramide levels and neuroinflammation, while levels of transcripts encoding proteins important for neuronal development and function were increased. Further, we show that nSMase2 deficiency reduces the level of exosomes transporting micro-RNAs that suppress expression of several/numerous genes important for neuronal function. These data indicate a critical role of nSMase2 in the regulation of cognitive decline during brain aging.

## Materials and Methods

### Animals and reagents

The *fro/fro* mouse was the gift from Dr. Christophe Poirier (Indiana University, Indianapolis). This carries a deletion of the C-terminal 33 amino acids in the sphingomyelin phosphodiesterase-3 (*Smpd3*) gene that encodes neutral sphingomyelinase 2 (nSMase2) (NCBI gene ID: 58994; (Aubin et al., 2005). Mice were handled according to the National Institutes of Health Guide for the Care and Use of Laboratory Animals. All procedures involving mice were approved by the Institutional Animal Care and Use Committee of University of Kentucky. Procedures focused on the use of male mice based on our previous studies showing that nSMase2 deficiency improves cognition in male 5XFAD mice (Dinkins et al., 2016). The antibodies, chemicals and reagents used in this study have been summarized and are listed in tables 2 and 3.

**Table 1.** List of Antibodies used in the study.

| **Antibodies** | **Ratio** | **Cat#** | **Company** |
| --- | --- | --- | --- |
| Anti-IL-1β | 1:250 | SC-32294 | Santa Cruz |
| Anti-IL-6 | 1:250 | SC-32296 | Santa Cruz |
| Anti-TNFα | 1:250 | SC-12744 | Santa Cruz |
| Anti-NR2B | 1:1000 | 06-600 | Millipore Sigma |
| Anti-Stat3 | 1:1000 | #4904 | Cell Signaling |
| Anti-pStat3 | 1:1000 | #9145 | Cell Signaling |
| Anti-P27 kip1 | 1:1000 | #2552 | Cell Signaling |
| Anti-ceramide rabbit IgG | 1:100 | Were generated in our laboratory |  |
| Anti-ceramide mouse IgM | 1:100 | MAS0014 | Glycobiotech |
| Anti-rabbit HRP | 1:5000 | 715-035-152 | Jackson Immune Research |
| Anti-mouse HRP | 1:5000 | 715-035-151 | Jackson Immune Research |
| Anti-GAPDH | 1:5000 | #5174 | Cell signaling tech |
| Anti-GFAP | 1:10,000 | 20071831 | DaKo |
| Anti-C3b | 1:1000 | 21337-1-AP | Protein Tech |

**Table 2.**
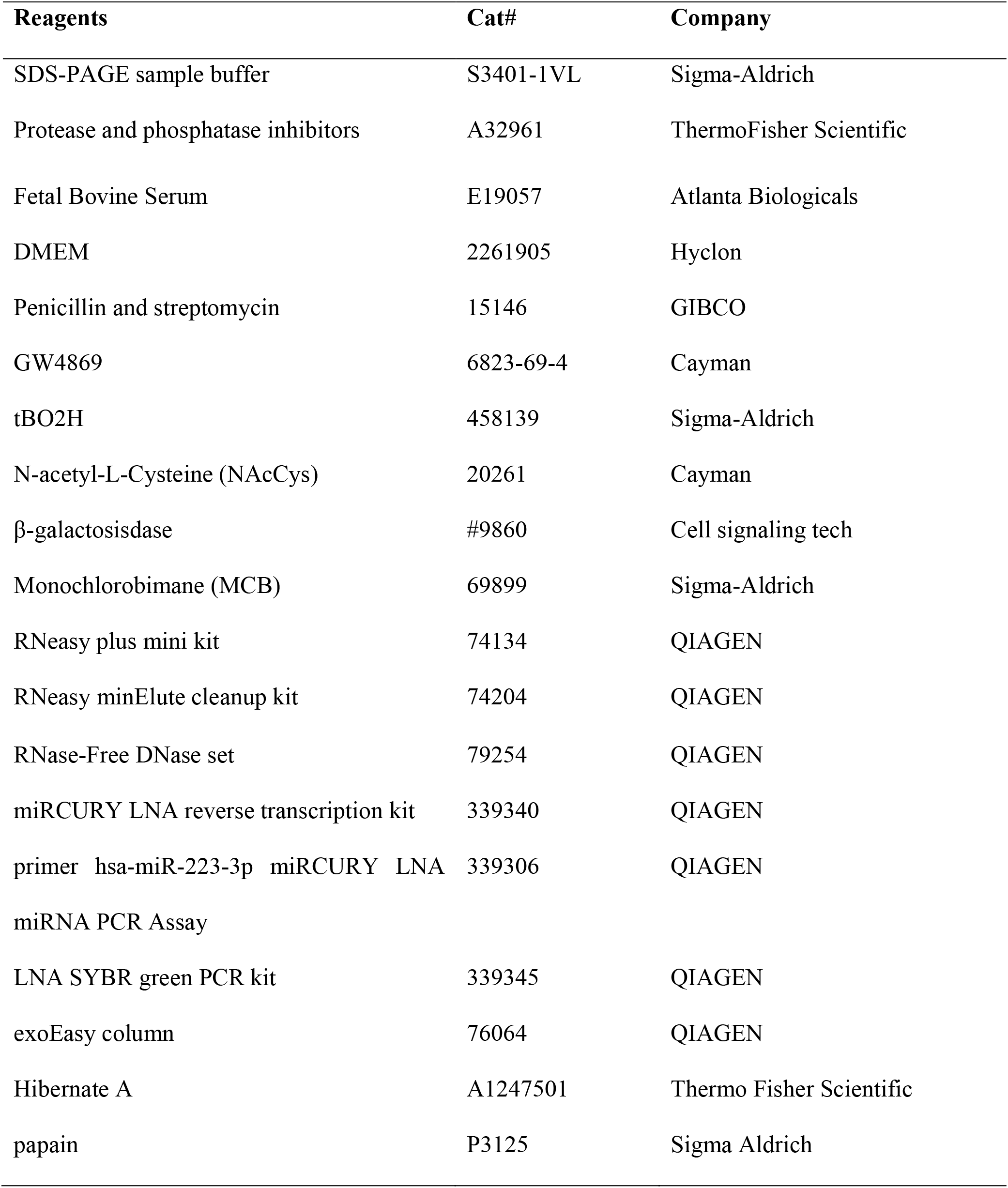
Chemicals and Kits used in the study.

**Table 3.** Positively behavior related diseases or Function Analysis.

| **Diseases or functions** | ***p*-value** | **Predicted** | **Activation** | **#Molecules** |
| --- | --- | --- | --- | --- |
| **Annotation** |  | **activation** | **z-score** |  |
|  |  | **state** |  |  |
| Cognition | 4.04E-23 | Increased | 2.995 | 118 |
| Learning | 1.76E-20 | Increased | 3.008 | 106 |
| Conditioning | 2.47E-09 | Increased | 2.976 | 46 |
| Memory | 1.26E-12 | Increased | 3.591 | 64 |

### Primary cell culture

Primary astrocytes from *fro/fro* and littermate controls (wild type or heterozygous +/*fro*) mice were prepared according to the protocol we have used previously (Dinkins et al., 2016; Kong et al., 2018). Astrocytes were maintained in DMEM with 10% FBS and 1% penicillin/streptomycin at 37 °C in a humidified atmosphere containing 5% CO2. For all treatment procedures, including incubation with reagents and isolation of extracellular vesicles, astrocytes were first maintained for 48 h in serum-and phenol-red free DMEM medium.

### Immunoblotting

Mouse brain cortex was solubilized by sonification in SDS-sample buffer containing 10% 2-mercaptoethanol and heated for 10 min at 95 °C prior to SDS-PAGE and immunoblotting. In detail, proteins were resolved by SDS gel electrophoresis on polyacrylamide gels (Biorad, Hercules, California) and transferred to nitrocellulose membrane (Bio-Rad, Hercules, California). Non-specific binding sites were blocked with 5% fat-free dry milk (Bio-Rad Hercules, California) in PBS containing 0.1% Tween-20 (PBST) followed by overnight incubation with primary antibodies. The primary and secondary antibodies diluted with different concentrations in PBST are listed in the table 1 of antibodies and reagents. Signals were detected using either pico or femto chemiluminescent (ECL) horseradish peroxidase (HRP) substrate (Thermo Fisher, Massachusetts, USA). Blot images were captured by Azure c600 system (Azure Biosystems, California, USA).

### Immunocytochemistry and fluorescence microscopy

Astrocytes grown on cover slips were fixed with 4% p-formaldehyde/0.5% glutaraldehyde/PBS for 20 min, followed by permeabilization with 0.2% Triton X-100 in PBS for 5-10 min at room temperature. Nonspecific binding sites were blocked with 3% ovalbumin/PBS for 1 hour at 37 °C. Cells were incubated with primary antibodies at 4 °C overnight. The next day, cells on cover slips were washed by PBS and followed by incubation with secondary antibodies for 2 h at room temperature. After washing with PBS, cover slips were mounted using Fluoroshield supplemented with DAPI (Sigma-Aldrich) to visualize the nuclei. Fluorescence microscopy was performed with a Nikon Ti2 Eclipse microscope equipped with NIS Elements software. Images were processed using a 3D deconvolution program as provided by the Elements software.

### Image analysis of colocalization studies

A set of images was de-identified (blinded identifiers) and numbers assigned using a random generator program (random.org) available online. Colocalization was analyzed using the Nikon Elements software. The degree of colocalization was assessed by calculation of the Pearson’s correlation coefficient for two fluorescence channels in overlays as previously described (Adler and Parmryd, 2010, 2013).

### Lipid analysis

For ceramide analysis, brain tissues were submitted to the lipidomics core facility at the Medical University of South Carolina, Charlston, SC (Dr. Besim Ogretmen, director) (https://hollingscancercenter.musc.edu/). Quantitative analyses of sphingolipids were based on previous published methods (Bielawski et al., 2009, 2010). The concentration of ceramide species was quantified in the sphingolipidomics (LC-MS/MS) analysis core. The lipid concentration was normalized to lipid phosphate.

### RNA-seq

RNA was extracted from the brain cortex using the miniRNeasy extraction kit (Qiagen). The total RNA was submitted to Novogene (https://en.novogene.com) for quality control and RNAseq analysis. Only the RNA with intergrity numbers ≥ 8, and of sufficient purity (OD260/280=1.8-2.2; OD260/230 ≥ 1.8) were used.

### Library preparation for transcriptome sequencing

A total amount of 0.4 ug of RNA was used for cDNA library construction at Novogene using an NEBNext® Ultra 2 RNA Library Prep Kit for Illumina® (cat NEB #E7775, New England Biolabs, Ipswich, MA, USA) according to the manufacturer’s protocol. Briefly, mRNA was enriched using oligo(dT) beads. Double-stranded complementary DNA was synthesized, beginning with priming by random hexamers. After terminal repair, poly-adenylation, and sequencing adaptor ligation, the cDNA libraries were size-selected and enriched by PCR. The resulting 250-350 bp insert libraries were quantified using a Qubit 2.0 fluorometer (Thermo Fisher Scientific, Waltham, MA, USA) and quantitative PCR. Libraries were sequenced on an Illumina NovaSeq 6000 Platform (Illumina, San Diego, CA, USA) using a paired-end 150 run (2×150 bases). The number of raw reads per sample ranged from 50 to 60 Million. Paired-end reads were aligned to the mouse mm10 build reference genome using the Spliced Transcripts Alignment to a Reference (STAR) software. Identification of differentially abundant transcripts was done using DEseq2. The resulting P values were adjusted using the Benjamini and Hochberg approach for controlling the false discovery rate, and the threshold for differential expression.

### Functional Bioinformatics analysis

The list of differentially abundant transcripts (*fro/fro vs +/fro*) was imported to the Kyoto Encyclopedia of Genes and Genomes (KEGG) database to identify pathway annotations enriched among the proteins encoded by these transcripts. The lists of differentially abundant transcripts (*fro/fro vs +/fro*) was also imported to the Ingenuity Pathway Analysis (IPA) software (Ingenuity H Systems, www.ingenuity.com)) to analyze functional annotations and regulatory networks via upstream regulator analysis, downstream effects analysis, mechanistic networks and causal network analysis.

### Exosome isolation and qRT-PCR

Exosomes were isolated from *+/fro* and *fro/fro* mice brain as described previously (Elsherbini et al., 2020b; Elsherbini et al., 2020a). miRNAs were eluted from exosomes using RNeasy minElute cleanup kit (Qiagen, 74204) and genomic DNA was digested with RNase-free DNase set (Qiagen, 79254). miRNA reverse transcription was performed using miRCURY LNA reverse transcription kit (Qiagen, 339340). qRT-PCR was performed to amplify microRNA 223-3p using the primer hsa-miR-223-3p miRCURY LNA miRNA PCR Assay (Qiagen, #339306, gene Globe ID: YP00205986) and LNA SYBR green PCR kit (Qiagen, 339345). Relative-fold changes were normalized comparing exosomal micro RNA reference gene miR-30c-5p using the primer hsa-miR-30c-5p miRCURY LNA miRNA PCR Assay (Qiagen 339306, gene globe ID YP00204783). Relative fold changes of miR223-3p of *fro/fro* mice were calculated to *+/fro* mice by using the formula: Relative value= 2^-Ct^geometrical mean of 223p-30c-5p and 30c-5p)^/2^Ct^geometrical mean of 30c-5p^. For determination of the amount of exosomes in *+/fro* and *fro/fro* mice, the right brain hemispheres of mice were digested in a mixture of papain (Sigma Aldrich, P3125) and hibernate A (Thermo Fisher Scientific, A1247501) at 37 °C for 20 min and centrifuged at 300×*g* for 5 min. Supernatants were centrifuged at 4000×*g* for 20 min and the resulting supernatants were recollected for further ultracentrifugation at 10,000×*g* for 40 min. Finally, exosomes were collected by passing the supernatants through exoEasy columns (Qiagen, 76064) and quantification was performed by nanoparticle tracking analysis (NTA) with ZetaView PMX110 (Particle Metrix).

### Statistical Analysis

Statistical analyses and graphing were performed using Microsoft Excel 2019 and GraphPad Prism 8.0 software (GraphPad, San Diego, CA, USA). For IPA analyses, a Z score (Z ≤-2.0 or Z ≥ 2.0) was considered significant. When the two groups were compared, student *t* test was used. Values of *P* <0.05 were considered significant. When multiple groups (>2) were compared, two-way ANOVA was applied.

## Results

### nSMase2 deficiency prevents age-related increase of ceramide levels in astrocytes

To determine if nSMase2 deficiency affects ceramide levels in the middle-aged brain, the cortex of 10-month old *fro/fro* and +/*fro* (heterozygous litermates) were subjected to targeted liquid chromatography-tandem mass spectrometry (LC-MS/MS). Figure 1 shows that the levels of all ceramide species were decreased in *fro/fro* cortex, with the ceramide species C_18:0_, C_18:1_, and C_24:1_ ceramide being downregulated the most. It was previously shown, that these ceramide species were upregulated in the aging brain and serum, and associated with memory impairment, including AD (Cutler et al., 2004; Haughey et al., 2010; Mielke et al., 2010a; Mielke et al., 2010b; Mielke and Lyketsos, 2010; Mielke et al., 2012).

**Figure 1.**
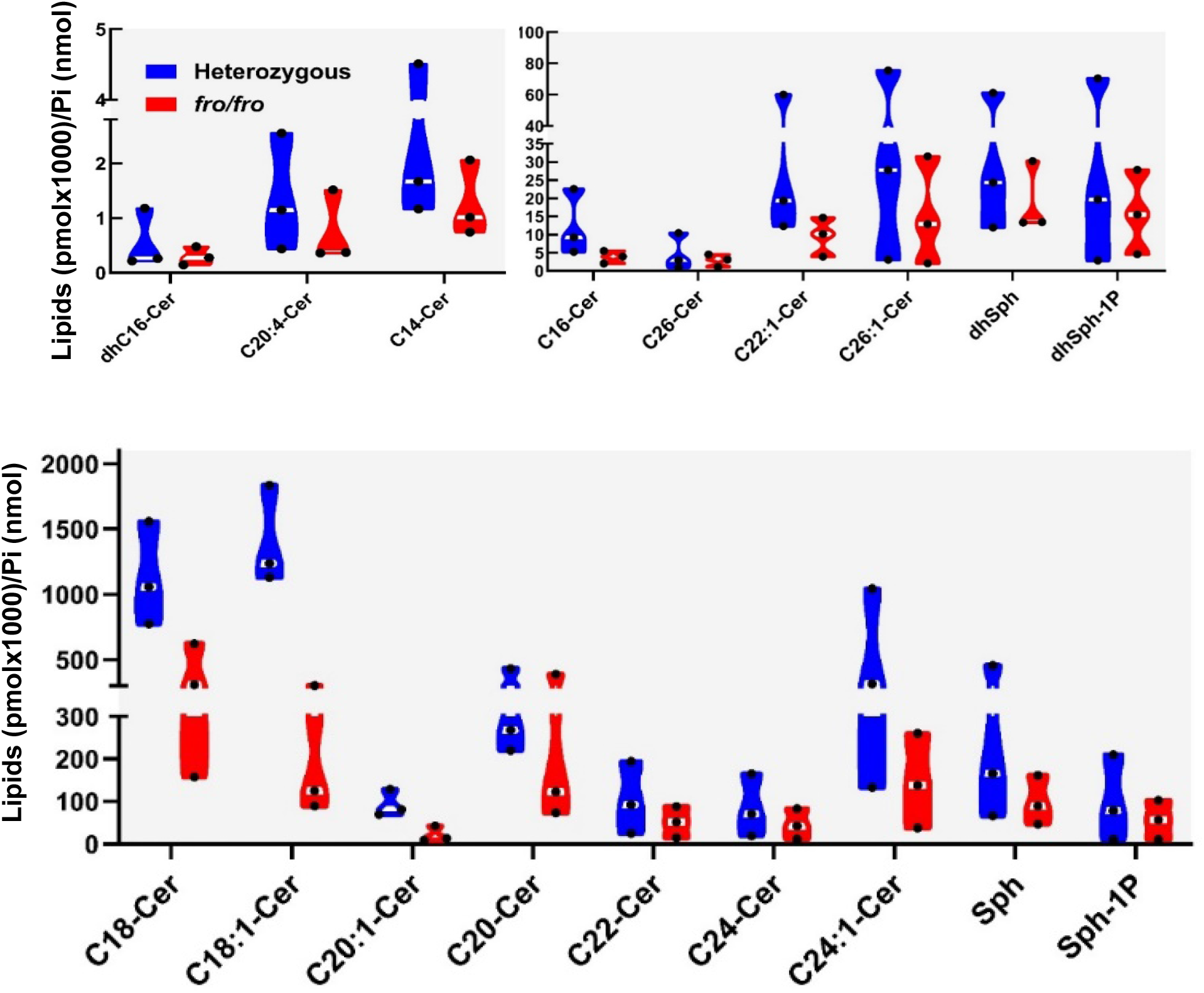
nSMase2 regulates the ceramide composition in the middle-aged brain. Sphingolipidomics (LC-MS/MS) analysis shows that the levels of all ceramide species are reduced in the nSMase2-deficient cortex (*fro/fro,* red) compared to heterozygous controls (+/*fro*, blue). Ceramide concentrations were normalized to lipid phosphate. N=3, 10-month old male, ratio-paired *t*-test; *p* <0.0001 for the difference in the levels of C24:1, C18:0, and C18:1 ceramide. Student’s *t-*test and ANOVA.

Using immunocytochemistry with an anti-ceramide antibody developed in our laboratory (Krishnamurthy et al., 2007), we tested if ceramide levels were affected depending on age and cell type. Figure 2 shows that labeling for ceramide was increased up to 4-fold in 10 and 15 month-old +/*fro* mice *vs*. 3 month-old mice, and in +/*fro* brains (from 15 month-old mice) when compared to 15 month-old *fro/fro* tissue. More importantly, increased ceramide labeling was predominantly colocalized with GFAP positive processes in *+/fro* tissue (arrows), while labeling in neurons was >2-times lower and confined to perinuclear compartments. In 10 and 15 month-old mice, punctate ceramide labeling was often juxtaposed to astrocytes indicating a secretory process (arrows in Fig. 2B-E). Ceramide labeling of microglia (Iba-1 positive) was not increased during brain aging and it was not altered by nSMase2 deficiency. These data suggest that nSMase2 deficiency mainly prevented elevation of ceramide and its secretion by astrocytes in the middle-aged brain.

**Figure 2.**
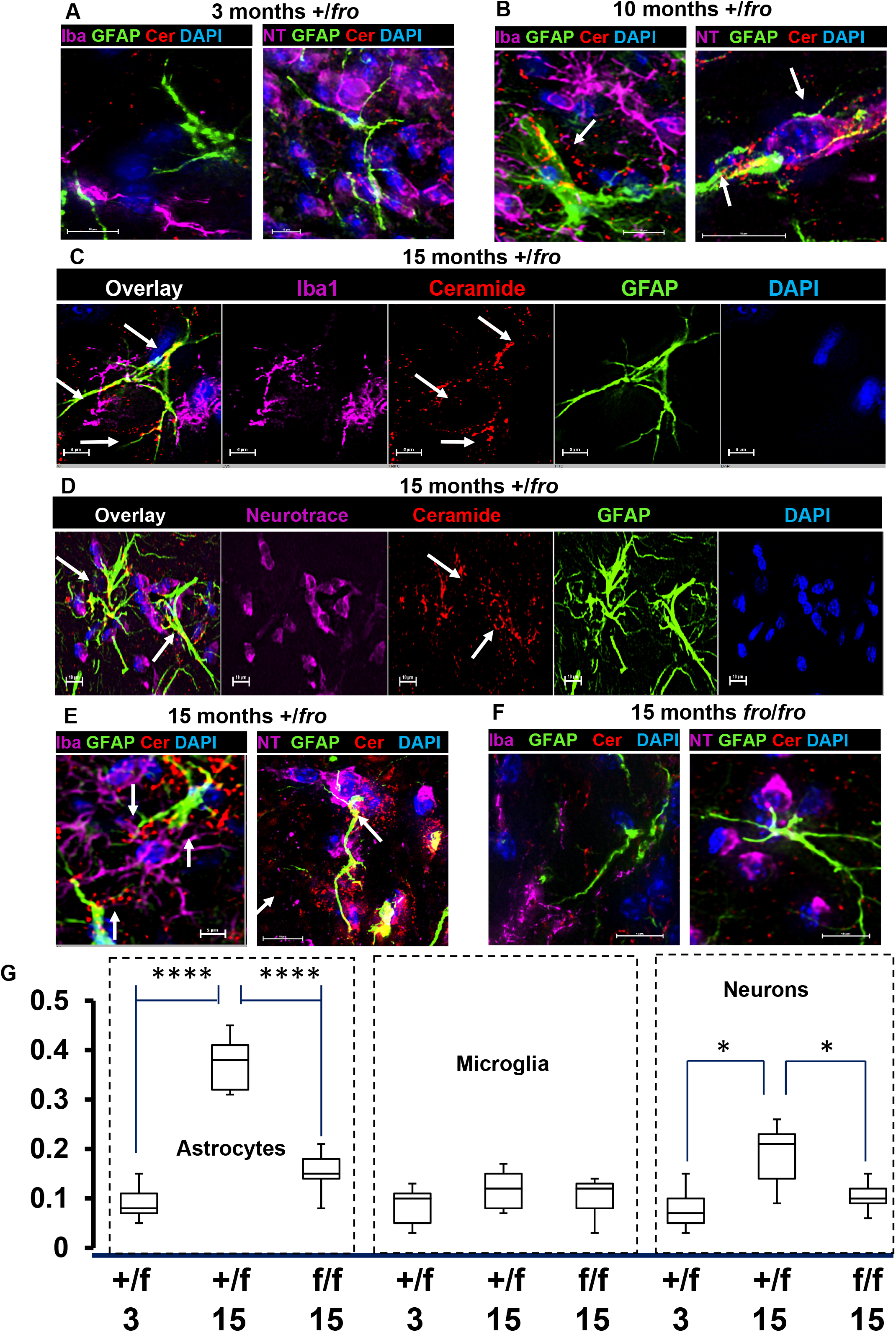
Ceramide is mainly increased in astrocytes during aging of the brain. (A-E) Immunocytochemistry shows that ceramide labeling (red) mainly increases in 10 and 15-month old (B-D, arrows) *vs*. 3-month old heterozygous (+/*fro*) cortex (A). Ceramide labeling is localized in astrocytes (GFAP, green) and to a lesser extent in microglia (purple in C) and neurons (purple in D). Punctate labeling of ceramide is also visible in close vicinity to astrocytic processes (arrows in B and E) and it is reduced in 15-month old *fro/fro* mice (E). (F) Quantitation of colocalization using Pearson’s coefficient for colabeling of ceramide with markers for astrocytes, (GFAP), microglia (Iba1), and neurons (Neurotrace). N=8. Unpaired *t*-test and ANOVA.

### nSMase2 deficiency alters mRNA abundance in the middle-aged brain

We performed RNAseq analyses on *+/fro* and *fro/fro* cortex to screen for clues as to how the lack of ceramide generation due to nSMase2 deficiency affects the middle-aged brain. A total of 1462 transcripts were changed, among them, 891 transcripts were increased and 571 transcripts were decreased. The most increased transcript was Grin2b (arrow in Fig. 3A). The differences in transcript abundance observed did not include differences in markers of specific cell types, and therefore did not argue for differences in the proportions of types of neurons or glia between the genotypes tested. This observation argues that these transcript abundance differences arose from differences in gene expression.

**Figure 3.**
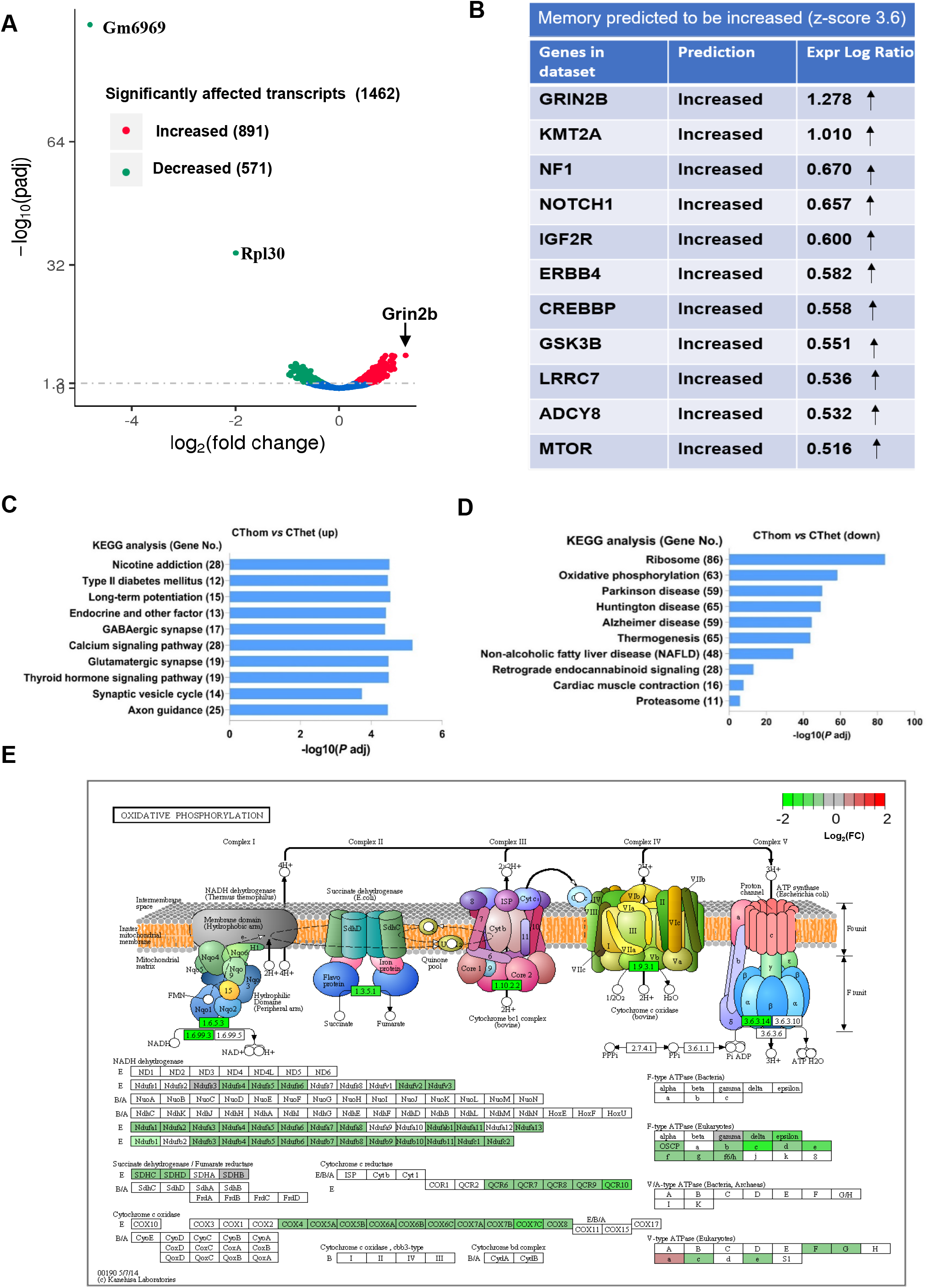
nSMase2-sensitive transcripts are related to oxidative phosphorylation, ribosome biogenesis, synaptic transmission, and memory. (A) Volcano plot shows differentially abundant transcripts in homozygous (*fro/fro*) vs. heterozygous (*+/fro*) mouse cortex. Red labels denote increased and green decreased transcripts in *fro/fro* cortex. The two transcripts decreased most are cytochrome c oxidase subunit VIIa polypeptide 2-like pseudogene (Gm6969) (and ribosomal protein L30 (RPL30) . The transcript most increased is Grin2b (NMDA receptor subunit 2B, arrow). (B) IPA analysis shows that Grin2b is one of a set transcripts encoding proteins important for memory that are increased in *fro/fro* mice. (C, D) KEGG pathway enrichment analysis of over-represented functional relationships among transcripts increased in *fro/fro* (CThom) cortex as compared to *+/fro* (CThet) cortex, showing the top 10 most significant biological pathways. (E) KEGG pathway analysis of oxidative phorphorylation shows the protein assembly in oxidative phosphorylation depicting five mitochondrial complexes containing proteins encoded by transcripts decreased in *fro/fro* mice. The color legend represents *fro/fro* vs. *+/fro* log2FC with red denoting upregulation, green denoting downregulation, and grey indicating log2FC is zero.

Functional bioinformatics analyses revealed that synaptic signaling annotations were over-represented among the transcripts that were increased in the nSMase2-deficient cortex and many of the encoded proteins, such as Grin2b, are important for memory (Fig. 3B and C). These analyses also revealed that nSMase2 deficiency was associated with increased levels of mRNAs encoding proteins involved in axonal guidance and long-term potentiation, suggesting that neural plasticity may be enhanced. Furthermore, these analyses revealed that annotations for neurodegenerative conditions such as AD, Parkinson’s disease, and Huntington’s disease were over-represented among the transcripts that decreased in *fro/fro* mouse brains (Fig. 3D). According to the KEGG analysis, also decreased was a set of transcripts encoding mitochondrial complex proteins important for oxidative phosphorylation, indicating that nSMase2 is involved in regulation of mitochondrial oxidative stress. (Fig. 3E). These discoveries are consistent with enhanced memory function and increased neural protection, and therefore provide the likely mechanistic explanations for the enhanced memory outcomes observed in *fro/fro* mice (Dinkins et al., 2016).

### nSMase2 deficiency or inhibition prevents oxidative stress-induced increase of ceramide levels and activation of astrocytes

In astrocytes, oxidative phosphorylation in mitochondria is reduced in favor of aerobic glycolysis, which was suggested to diminish oxidative stress and protect neurons (Demetrius and Simon, 2012; Tavallaie et al., 2020; Zheng et al., 2021). Oxidative stress, however, is increased in reactive astrocytes and induces the production of proinflammatory cytokines (Lee et al., 2010). Based on the KEGG pathway enrichment analysis showing that a set of transcripts encoding proteins participating in mitochondrial oxidative phosphorylation were decreased in *fro/fro* brain (Fig. 3D and E), we hypothesized that nSMase2 deficiency decreased astrocyte activation, which was consistent with reduced levels of transcripts for S100β and C1q, two markers for activated astrocytes (not shown).

Since astrocyte activation has been associated with increased oxidative stress and activation of nSMase2, we tested the response of primary cultured heterozygous (+/*fro*) and *fro/fro* astrocytes to oxidative stress induced by incubation with tertiary butyl peroxide (tBO2H). We used monochlorobimane (MCB), a fluorescence indicator for glutathione (GSH), and the anti-ceramide antibody to detect the level of GSH and ceramide in tBO2H-treated astrocytes using immunofluorescence microscopy. Figure 4A shows that incubation of astrocytes with tBO2H led to the reduction of MCB fluorescence in both, +/*fro* and *fro/fro* astrocytes, while ceramide labeling was only increased in +/*fro* astrocytes. This result indicated that oxidant-induced loss of GSH activated nSMase2 and led to the generation of ceramide in heterozygous astrocytes, which was absent in *fro/fro* astrocytes. Likewise, GFAP labeling, as a marker of activated astrocytes, was only increased in heterozygous cells, indicating that tBO2H incubation led to activation of astrocytes, which required activation of nSMase2 by oxidative stress. This conclusion was further supported by colocalization analyses showing that in control (non-treated) as well as tBO2H-treated heterozygous astrocytes, loss of MCB labeling was associated with strong ceramide labeling, and that the increase of ceramide labeling was directly correlated with that of GFAP (Pearson’s colocalization coefficient 0.65±0.15).

**Figure 4.**
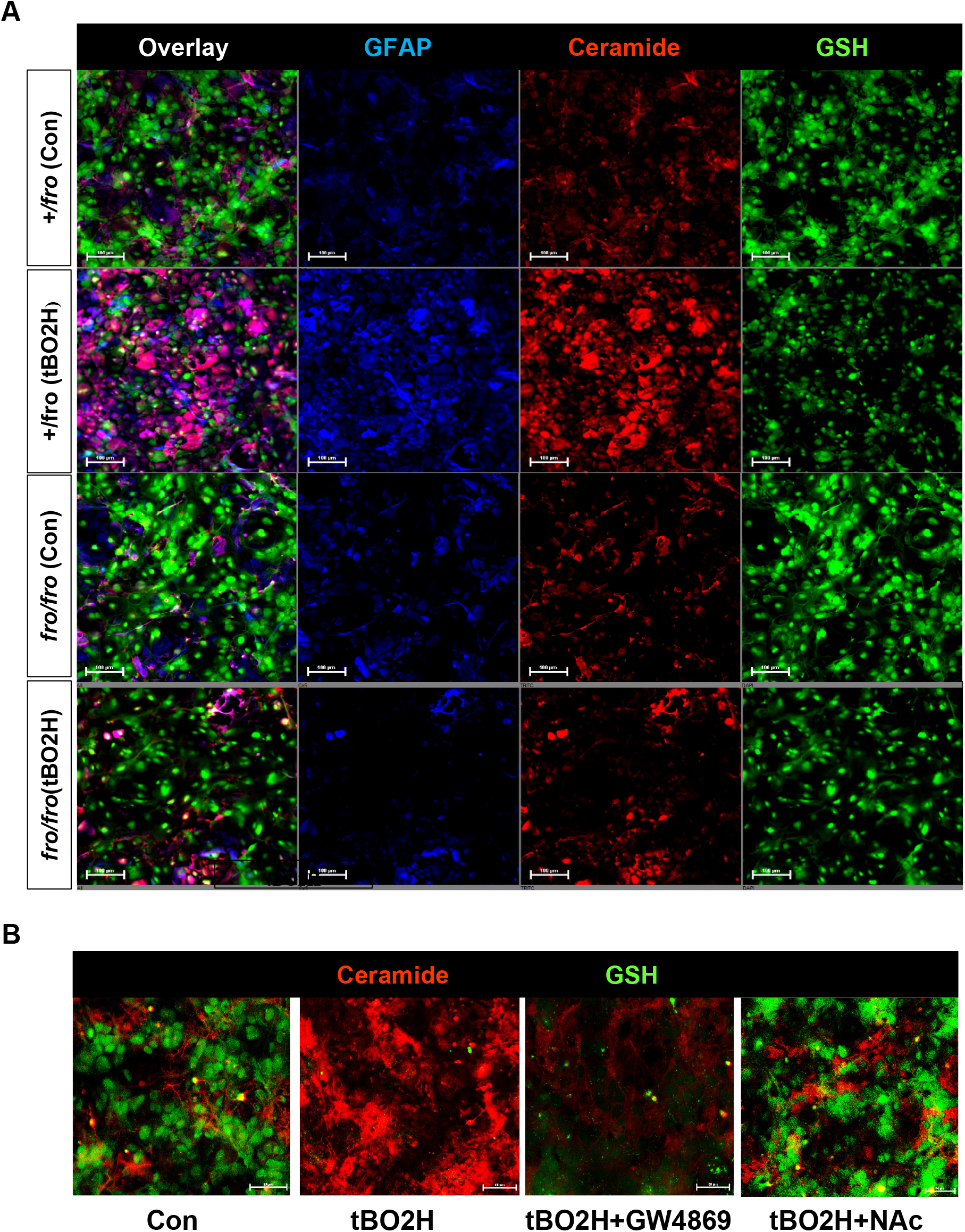
Oxidative stress reduces GSH levels and increases nSMase2-mediated ceramide generation in reactive astrocytes. (A) Oxidative stress induced by treatment of heterozygous (*+/fro*) astrocytes for 1 h with 200 µM tBO2H led to the depletion of GSH (MCB labeling) and increase of ceramide and GFAP labeling, while nSMase2-deficient (*fro/fro*) astrocytes showed no increase of ceramide or GFAP levels. There is no colabeling of GSH with ceramide. (B) Inhibition of nSMase2 with GW4869 (10 µM) or restoration of GSH with NAcCys (500 µM) prevents ceramide generation induced by oxidative stress (100 µM tBO2H for 2 h).

To test if restoration of GSH levels prevented oxidative-stress induced activation of nSMase2 and generation of ceramide, we pre-incubated tBO2H-treated wild type astrocytes with N-acetyl cysteine (NAcCys), an endogenous antioxidant known to provide cysteine to boost GSH production and reduce oxidative stress *in vitro* and *in vivo* (Farr et al., 2003; Rutkute et al., 2007; Cao et al., 2012; Skvarc et al., 2017). Figure 4B shows that NAcCys (NAc) prevented depletion of GSH and increase of ceramide labeling. GW4869, an inhibitor of nSMase2, did not block tBO2H-induced GSH depletion, but prevented the increase of ceramide in the presence of tBO2H, confirming that onset of oxidative stress is upstream of nSMase-2 activation and generation of ceramide. Taken together, these results suggest that oxidative stress leads to the decrease of GSH levels and nSMase2-mediated generation of ceramide.

### nSMase2 deficiency reduces astrocyte senescence and inflammation in the brain

Oxidative stress-induced astrocyte activation has been associated with inflammation and astrocyte senescence in the aging brain (Lee et al., 2010). Further, the onset of senescence in astrocytes is an essential part of the mechanisms underlying functional decline during brain aging (Cohen and Torres, 2019). Since both, oxidative stress and ceramide have been implicated in cellular senescence, we tested if nSMase2 deficiency delayed senescence of astrocytes. We assayed the activity of a senescence marker, β-galactosidase (β-gal), in 5-weeks old primary cultures of +/*fro vs*. *fro/fro* astrocytes. Figure 5A and B shows that β-gal staining was lower in nSMase2-deficient astrocytes (*fro/fro)* as compared to +/*fro* astrocytes, even after normalizing for cell density suggesting that lack of nSMase2 delayed aging of astrocytes. β-gal staining in aging +/*fro* astrocytes was similar to that in wild type astrocytes, indicating that a single allele of nSMase2 is sufficient for ceramide generation underlying astrocyte senescence (not shown).

**Figure 5.**
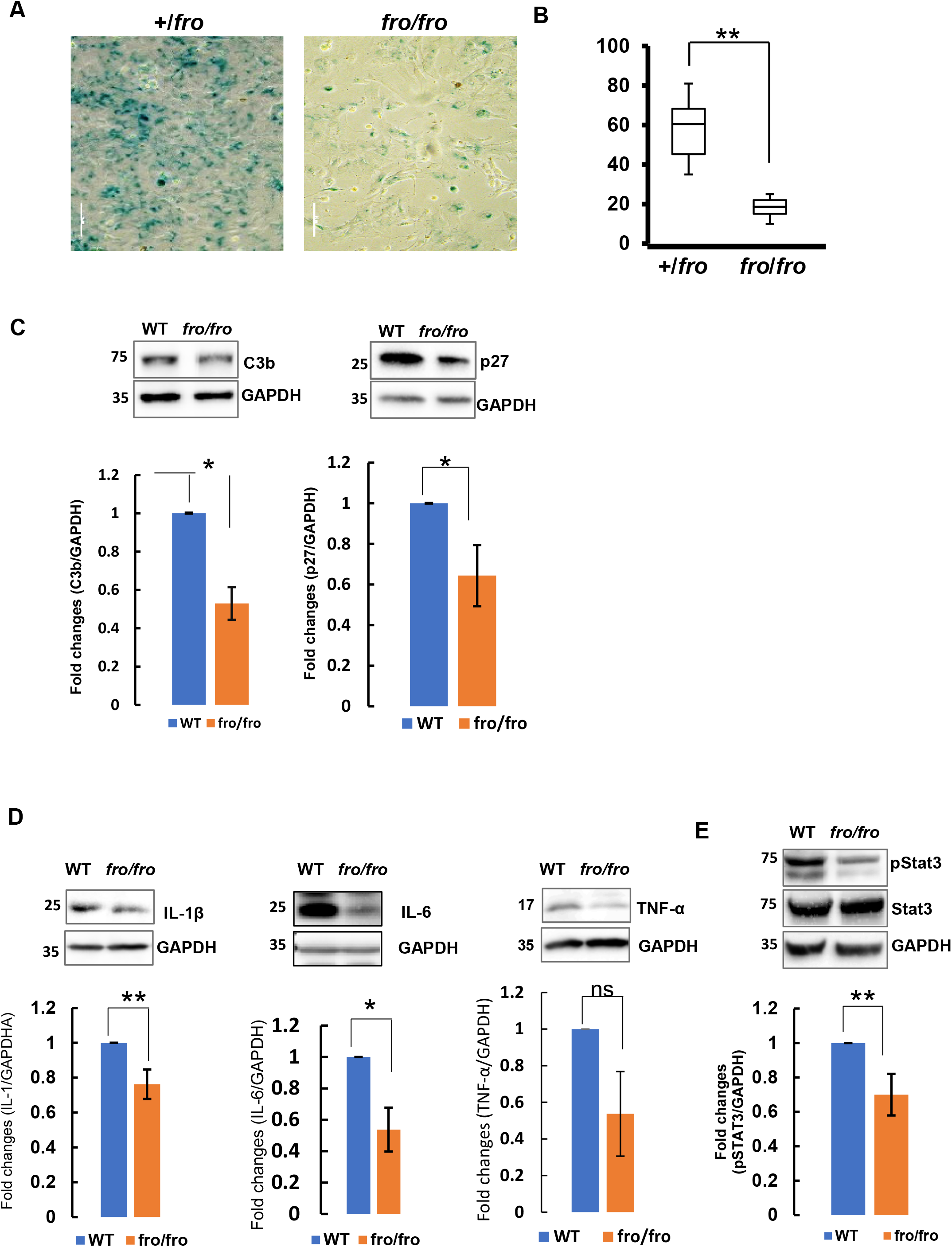
nSMase2 regulates expression of senescence markers, pro-inflammatory cytokines and Stat3 activation. (A, B) Five-weeks old cultures of primary *fro/fro* astroctyes show significantly less β-galactosidase staining than heterozygous (+/*fro*) astrocytes. N=3. (C-E) Immunoblots for senescence markers p27 and C3b (C), cytokines (IL-1β, IL-6, and TNF-α) (D), and pStat3 and Stat3 (E) in protein prepared from 10-months old male wild type (WT) and nSMase2-deficient (*fro/fro*) cortex. N=3. Unpaired *t*-test.

Based on the results of the β-gal *in vitro* assay, suggesting that senescence was decreased in nSMase2-deficient astrocytes, we determined the level of senescence markers in *fro/fro* brain tissue. Consistent with the *in vitro* data in primary astrocytes, Figure 5C shows that the levels of two senescence markers, p27 and C3b were lower in the middle-aged *fro/fro* brain as compared to the wild type control. Since senescence in the brain is associated with neuroinflammation, we next determined the level of proinflammatory cytokines using immunoblot analysis. Figure 5D shows that IL-1β and IL-6, were significantly decreased in *fro/fro* mouse brain, suggesting that nSMase2 deficiency reduces inflammation in the middle-aged brain.

Upregulation of proinflammatory or senescence markers is characteristic of reactive astrocytes (Escartin et al., 2021). Proinflammatory cytokines induce tyrosine (Y705) phosphorylation of Stat3, a transcription factor activated in reactive and aging astrocytes (Herrmann et al., 2008; Hashioka et al., 2011; O’Callaghan et al., 2014; White et al., 2020). Consistent with decreased levels of proinflammatory cytokines, we observed significantly reduced phosphorylation of Stat3 in *fro/fro* brains as compared to wild type (Fig. 5E). This indicates that nSMase2 participates in the upregulation of inflammation in the middle-aged brain through activation of the Stat3 signaling pathway.

### nSMase2 deficiency increases axonal growth and glutamate signaling transcripts

In contrast to the suppressive effect of nSMase2 deficiency on several transcripts encoding proteins important in astrocyte activation, senescence, and neuroinflammation, our data clearly show increases in sets of transcripts encoding proteins involved in axonal guidance/growth and neural signal transmission, especially synaptic signaling and plasticity related to cognition, memory, and learning (Table 3). Figure 6A displays the primary network identified when the whole set of differentially abundant transcripts was analyzed by IPA. The sub-networks include motor-function related coordination, processing of RNA and several transcription regulation networks, such as those driven by MYC and MYCN. The motor-related behavioral coordination networks involve Grin2b, Grin2a, and BDNF. BDNF is predicted to positively regulate Grin2b in coordination-associated networks (Fig. 6A, B) (Zhang et al., 2018). More importantly, Grin2b is a critical component of the N-methyl-D-aspartate receptors that mediate forms of long-term potentiation fundamental to cognitive functions and memory (Fig. 7A) (Gozlan et al., 1995). To confirm that upregulation of Grin2b transcript results in increased amount of protein, we determined the protein levels of Grin2b in the brain cortex of both wild type and *fro/fro* mice (10-month old). Consistent with the RNAseq data, we found that the protein level of Grin2b in the brain cortex was two-times higher in *fro/fro* mice than in wild type mice (Fig. 7B).

**Figure 6.**
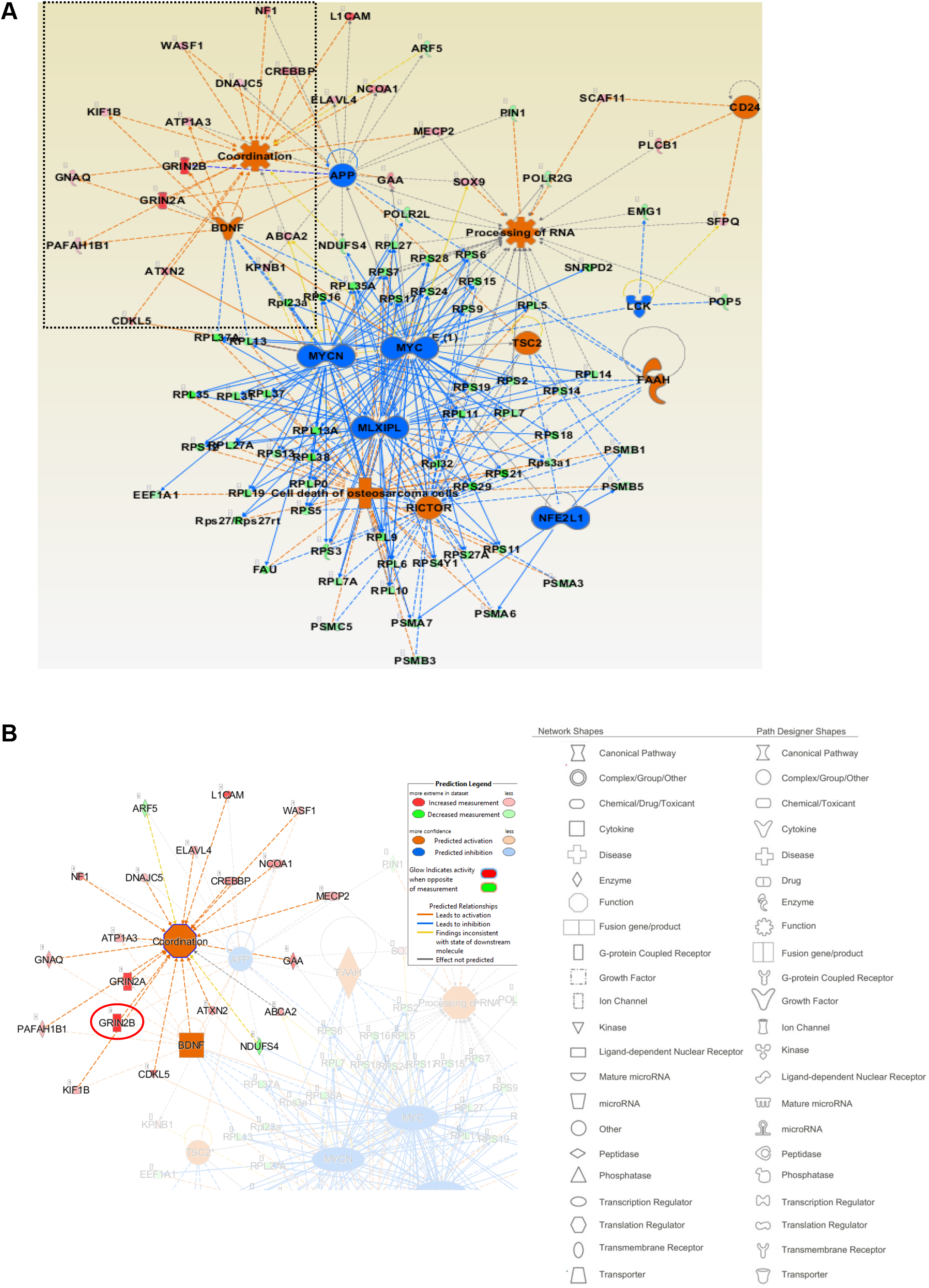
nSMase2 regulates Grin2b transcription networks. (A) IPA analysis of the set of differentially abundant transcripts shows the most enriched mechanistic network, including motor behavioral coordination-associated networks and processing of RNA. The legends of network and path designer shapes have been listed underneath of networks, red color denotes activated function and blue color denotes inhibited function. (B) Grin2b, Grin2a, and BDNF were predicted to be coordinatedly upregulated in *fro/fro* cortex.

**Figure 7.**
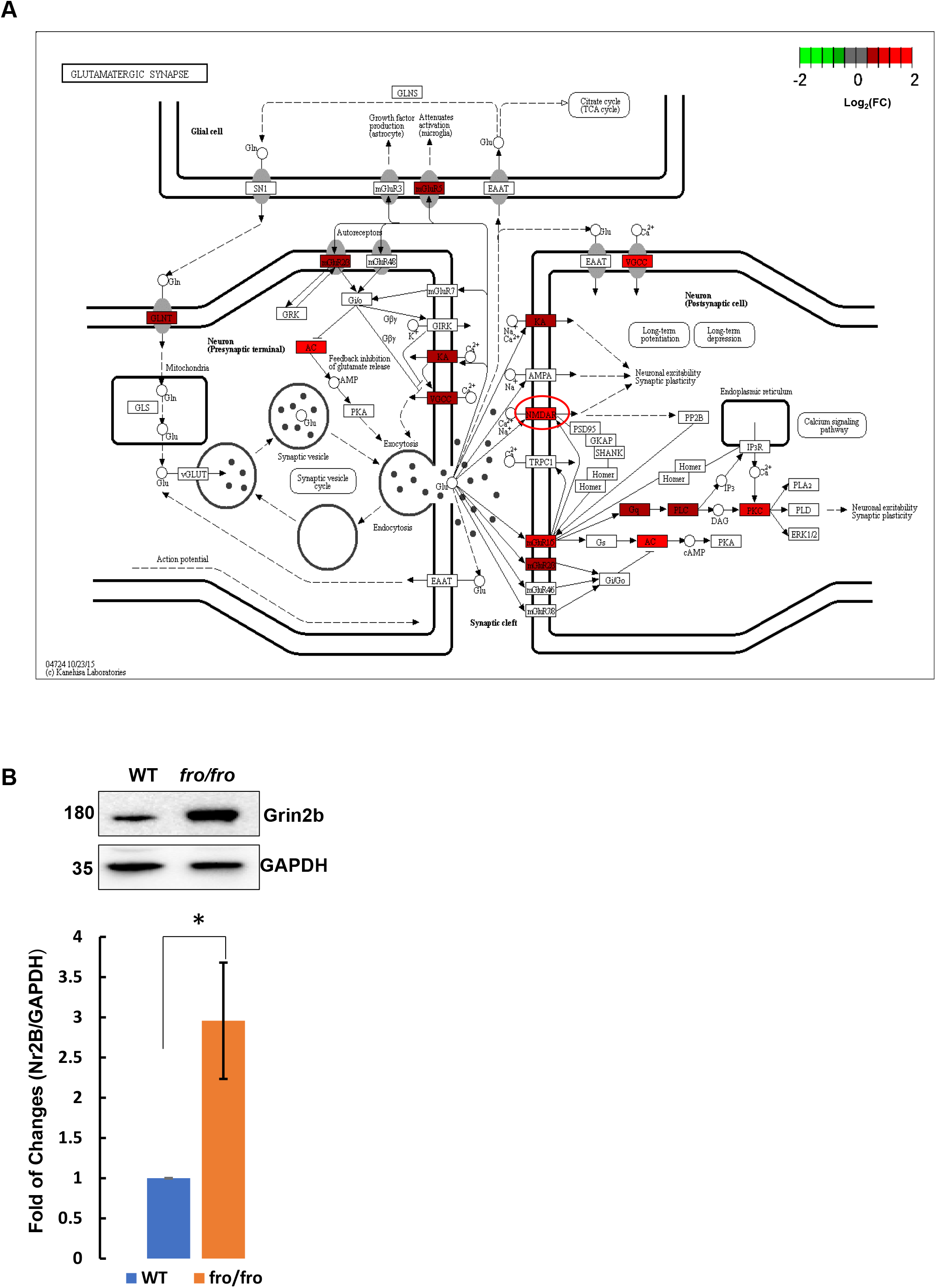
nSMase2 deficiency increases Grin2b expression. (A) Synaptic network for receptor cell signaling upregulated in *fro/fro* cortex, including NMDA receptor (circled in red) containing Grin2a and Grin2b subunits. (B) Protein analysis of Grin2b in *fro/fro* cortex as compared to wild type controls. N=3 pairs (littermates from 3 dams), two tails paired *t*-test.

Increases in Grin2b mRNA may be induced by nSMase2 deficiency in neurons or other cell types, such as astrocytes. Since immunolabeling for ceramide was increased in astrocytes of the aging brain, we investigated a potential trans-cellular effect of astrocytes on gene expression in neurons. Two potential factors suppressing expression of Grin2b are the neural differentiation transcription factor REST and the micro-RNA miR-223-3p (Harraz et al., 2012; Rodenas-Ruano et al., 2012) (Fig. 8A). We previously published that reactive astrocytes secrete exosomes enriched with ceramide, consistent with punctate labeling for ceramide in juxtaposition to astrocytic processes in 10 and 15-month old mouse cortex (Fig. 2) (Wang et al., 2012). A recent study showed that exosomes secreted by reactive astrocytes were enriched with miR-223-3p and downregulated Grin2b mRNA when added to neuronal cultures (Amoah et al., 2020). To test if nSMase2 regulated the amount of exosomal miR-223-3p, we performed qRT-PCR for miR-223-3p in brain derived-exosomes from 10-month old *+/fro* and *fro/fro* mice. This experiment did not reveal a significant decrease when compared to exosomal housekeeping micro-RNA (Fig. 8B), but it also did not rule out contributions from small changes in miR-223-3p to the increased amounts of Grin2b in *fro/fro* mice. More importantly, quantitation of exosomes showed that their number was reduced by 70% in the *fro/fro* cortex when compared to that of +/*fro* mice (Fig. 8C and D). Therefore, even if the copy number/exosomes of miR-223-3p is not significantly reduced, the decrease of exosomes secretion due to nSMase2 deficiency will lower the impact of this micro-RNA on the amount of Grin2b in *fro/fro* mice.

**Figure 8.**
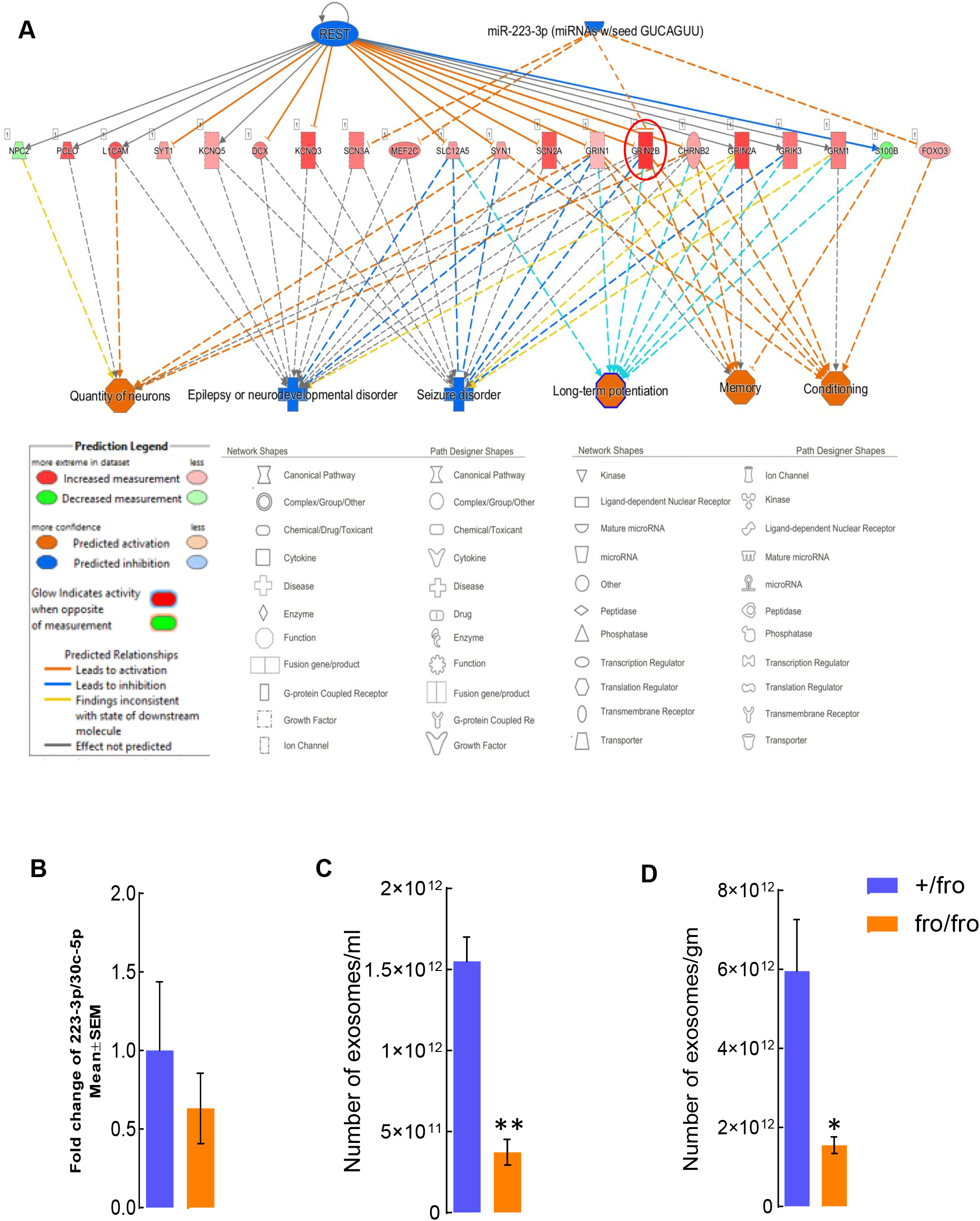
nSMase2 deficiency reduces the number of exosomes that decrease Grin2b. (A) The upstream regulator analysis by IPA showing transcripts downstream of REST and miR-223-3p that decreased in *fro/fro* cortex. The inhibited function of REST and miR-223-3p as shown in blue color positively activate downstream Grin2b. The transcripts downstream of REST and miR-223-3p are positively associated with neuron number, long-term potentiation, epilepsy, behavioral conditioning, and memory. (B) qPCR for miR-223-3p in exosomes from *fro/fro* and *+/fro* brain. Plot shows fold-change when compared to exosomal house keeping miRNA (miR-30c-5p). N=3, unpaired *t*-test. (C, D) Nanoparticle tracking analysis (Zetaview) of extracellular vesicles from +/*fro* and *fro/fro* cortex. The mumber of exosomes are shown as per militer (ml) of PBS or per gram (gm) of brain tissues N=3. Unpaired *t*-test.

## Discussion

We have previously shown that nSMase2 deficiency (*fro/fro* mouse) led to improved cognition in 5XFAD mice (Dinkins et al., 2016). During these studies, we detected marked improvement of memory function (fear conditioning test) in non-AD *fro/fro* controls when compared to wild type littermates, suggesting that reduced cognitive decline due to nSMase2 deficiency involves mechanisms independent of AD. To identify these mechanisms regulated by nSMase2, we performed transcriptomic (RNAseq) analyses on cortices of homozygous (*fro/fro*) and heterozygous (+/*fro*) mice. We focused on middle-aged male mice, since our studies on AD showed that cognitive improvement due to nSMase2 deficiency was limited to middle-aged males.

Consistent with improved cognition, RNA-seq analysis revealed increases in clusters of transcripts encoding axonal guidance and synaptic signal transmission proteins in the cortex of *fro/fro* mice (Table 3, Fig. 3). The coordinated increase in these sets of functionally related transcripts due to nSMase2 deficiency suggests that nSMase2 regulates neuronal function either intrinsically or by trans-cellular effects arising from other cell types, particularly astrocytes. Evidence against intrinsic effects arises from decreases in transcripts encoding mitochondrial proteins important for oxidative phosphorylation in *fro/fro* cortex (Fig. 3). Oxidative phosphorylation is critical for neurons, but not for astrocytes, which mainly generate ATP from aerobic glycolysis (Demetrius and Simon, 2012; Zheng et al., 2021). Hence, if our evidence for decreased capacity for oxidative phosphorylation stems from downregulation of oxidative phosphorylation in neurons this would impede enhanced synaptic transmission and cognitive performance. In addition, recent studies show that downregulation of oxidative phosphorylation and upregulation of aerobic glycolysis in astrocytes is protective for neurons and ameliorates AD pathology (Demetrius and Simon, 2012; Zheng et al., 2021). Therefore, the current study is based on the hypothesis that decreases in transcripts encoding proteins involved in oxidative phosphorylation are limited to astrocytes. Indeed, the results are consistent with this hypothesis showing reduction of glial activation and neuroinflammation as indicated by decreased levels of S100β and C1q transcripts and proinflammatory cytokines (Fig. 3 and 5D), as well as reduced astrocyte senescence (Fig. 5A and B).

Increased astrocyte activation and neuroinflammation are hallmarks of the aging brain and associated with reduced levels of GSH, an endogenous antioxidant mainly produced in astrocytes. There is a strong correlation between cognitive decline and lower levels of GSH (Hajjar et al., 2018). The GSH levels in the brain decrease with age, mainly by being used to neutralize reactive oxygen species (ROS) (Zhu et al., 2006). Astrocytes are the main source for the GSH precursor dipeptide (Cys-Gly) used for GSH biosynthesis in neurons (McBean, 2017). Astrocytes maintain their neuroprotective role under acute oxidative stress (Bhatia et al., 2019), but they become depleted of GSH during aging and neurodegenerative disease (Lee et al., 2010). The immediate target affected by GSH depletion in astrocytes is still unknown. Our *in vitro* experiments with primary cultured astrocytes show that reduction of GSH due to oxidative stress induces generation of ceramide, which is prevented by inhibition or deficiency of nSMase2 (Fig. 4). It is known that the GSH concentration found in astrocytes (5-8 mM) is in the range required for inhibition of nSMase2 (Liu and Hannun, 1997; Liu et al., 1998; Rutkute et al., 2007; Nikolova-Karakashian et al., 2008; Lee et al., 2010; McBean, 2017). Hence, oxidative stress leading to GSH depletion will activate nSMase2 and in turn, induce generation of ceramide.

Lipidomics analysis shows that in the aging brain, all major ceramide species, including those found to be predominantly reduced in nSMase2-deficient *fro/fro* brain (C18:0, C18:1, and C24:1 ceramides) are elevated (Cutler et al., 2004; Haughey et al., 2010; Mielke et al., 2010a; Mielke et al., 2010b; Mielke and Lyketsos, 2010; Mielke et al., 2012). Since the reduction of GSH levels in the aging brain is strongly associated with cognitive decline, we explored the hypothesis that the decrease of GSH levels leads to the activation of nSMase2 and generation of ceramide during aging. The strongest driver of GSH depletion during aging is oxidative stress leading to reactive oxygen metabolites that deplete GSH. Therefore, it is reasonable to conclude that the *in vitro* experiments showing increased generation of ceramide by oxidative stress-induced activation of nSMase2 in astrocytes recapitulate a mechanism similar to that in the aging brain.

Previous studies have shown that oxdative stress activates astrocytes and leads to the secretion of proinflammatory cytokines (Escartin et al., 2021). Since these cytokines induce activation of nSMase2 and ceramide participates in mediating signaling downstream of cytokine receptors, we tested if neuroinflammation is reduced in nSMase2-deficient brain. The immunoblot data on inflammation and senescence markers showed that *fro/fro* mice cortex contained decreased levels of the proinflammatory cytokines IL-1β, IL-6, and TNF-α, and lower levels of the senescence markers p27 and C3b (Fig. 5C and D). These data are consistent with a lower phosphorylation level of Stat3 (Y705), the major protein kinase activated by proinflammatory cytokines, upregulated during aging, and characteristic for reactive astrocytes (Fig. 5E) (Herrmann et al., 2008; Hashioka et al., 2011; O’Callaghan et al., 2014; White et al., 2020). Downregulation of the Stat3-associated cell signaling pathways suggests that aging-related processes, particularly activation and senescence of astrocytes are prevented or delayed in nSMase2-deficient *fro/fro* brain. This result is consistent with our *in vitro* data on nSMase2-deficient astrocytes, which age slower than wild type astrocytes as shown by β-galactosidase staining in 5-week old primary cultures of *fro/fro vs*. +/*fro* astrocytes (Fig. 5A and B).

In agreement with delayed brain aging, nSMase2 deficiency is associated with sets of transcripts that encode proteins involved in axon development and synaptic signal transmission (Fig. 3). Particularly, the mRNA and protein levels of Grin2b, the b-subunit of the ionotropic glutamate receptor are increased by about 2-fold in *fro/fro* cortex (Figs. 3A and 7B). The level of Grin2b upregulation is similar to that of a transgenic mouse generated about 20 years ago (Tang et al., 1999; Cao et al., 2007). The aged Grin2b overexpressing mice outperformed their wild-type littermates in five different learning and memory behavior tests, including fear conditioning tests. In fact, the performance of these mice in the fear conditioning test was very similar to that of the *fro/fro* mouse, consistent with a similar expression level of Grin2b. Conversely, the subunit composition of the NMDA receptor shows a significant decrease in NR1, NR2A, and NR2B (Grin2b) at the synaptic protein level during aging and in AD (Avila et al., 2017). This raises the interesting possibility that downregulation of the NMDA receptor, especially the Grin2b subunit, contributes to decreased cognitive function during the aging process. Therefore, the increased level of Grin2b in *fro/fro* mice suggests that nSMase2 deficiency is beneficial for learning and memory function in the middle-aged brain.

Since nSMase2 deficiency decreases the level of ceramide, it is critical to understand how and in which cell type lack of nSMase2 activity prevents ceramide-induced aging of the brain and cognitive decline. Since nSMase2 deficiency is not confined to astrocytes, reduction of ceramide levels may be critical in neurons and other cells as well. Immunolabeling for ceramide in cryosections shows that the level of ceramide is mainly increased in astrocytes of the aging brain, and lower in astrocytic processes of *fro/fro* brain (Fig. 2). In addition, punctate labeling of ceramide in close vicinity to GFAP (+) astrocytic processes indicates ceramide secretion, which is known for reactive astrocytes that secrete ceramide-enriched exosomes or “astrosomes” (Wang et al., 2012). These data suggest that aging-related increase of ceramide in wild type astrocytes may have a trans-cellular effect on other cells such as neurons via secretion of exosomes.

Recently, it was shown that activated astrocytes secrete exosomes that are enriched with miR-223-3p, a micro-RNA that downregulates Grin2b levels in neurons (Fig. 8A) (Amoah et al., 2020). Our data show reduced levels of exosomes in *fro/fro* brain, including those carrying miR-223-3p (Fig. 8B-D). These data are consistent with our previous studies showing that nSMase2 deficiency or inhibition reduced exosome secretion from *in vitro*-cultivated astrocytes and brain exosome levels (Wang et al., 2012; Dinkins et al., 2016). Therefore, it is possible that nSMase2 deficiency reduces exosomal micro-RNAs that downregulate neuronal function, which could explain the increase of the respective mRNAs levels in *fro/fro* brain.

One may speculate that GSH and ceramide metabolism are intertwined in astrocytes and neurons of the aging brain (Fig. 9). In astrocytes, oxidative phosphorylation is downregulated in favor of aerobic glycolysis, which produces reduction equivalents (NADH) to regenerate GSH. GSH inhibits nSMase2-mediated ceramide generation and secretion of astrosomes. Increased oxidative stress during aging leads to reduction of GSH levels in astrocytes and activation of nSMase2, which induces formation and secretion of aging-associated astrosomes (AAAs) that suppress neuronal activity. In nSMase2-deficient (*fro/fro*) mice, lack of ceramide elevation prevents formation of AAAs, and in turn, preserves neuronal function. Hence, GSH as a sensor for oxidative stress and nSMase2-catalyzed generation of ceramide may constitute a feedback loop for the function of AAAs in the regulation of neuronal activity. In our future studies, we will further investigate the role of nSMase2 and ceramide in oxidative stress and aging-related regulation of neuronal activity and cognition.

**Figure 9.**
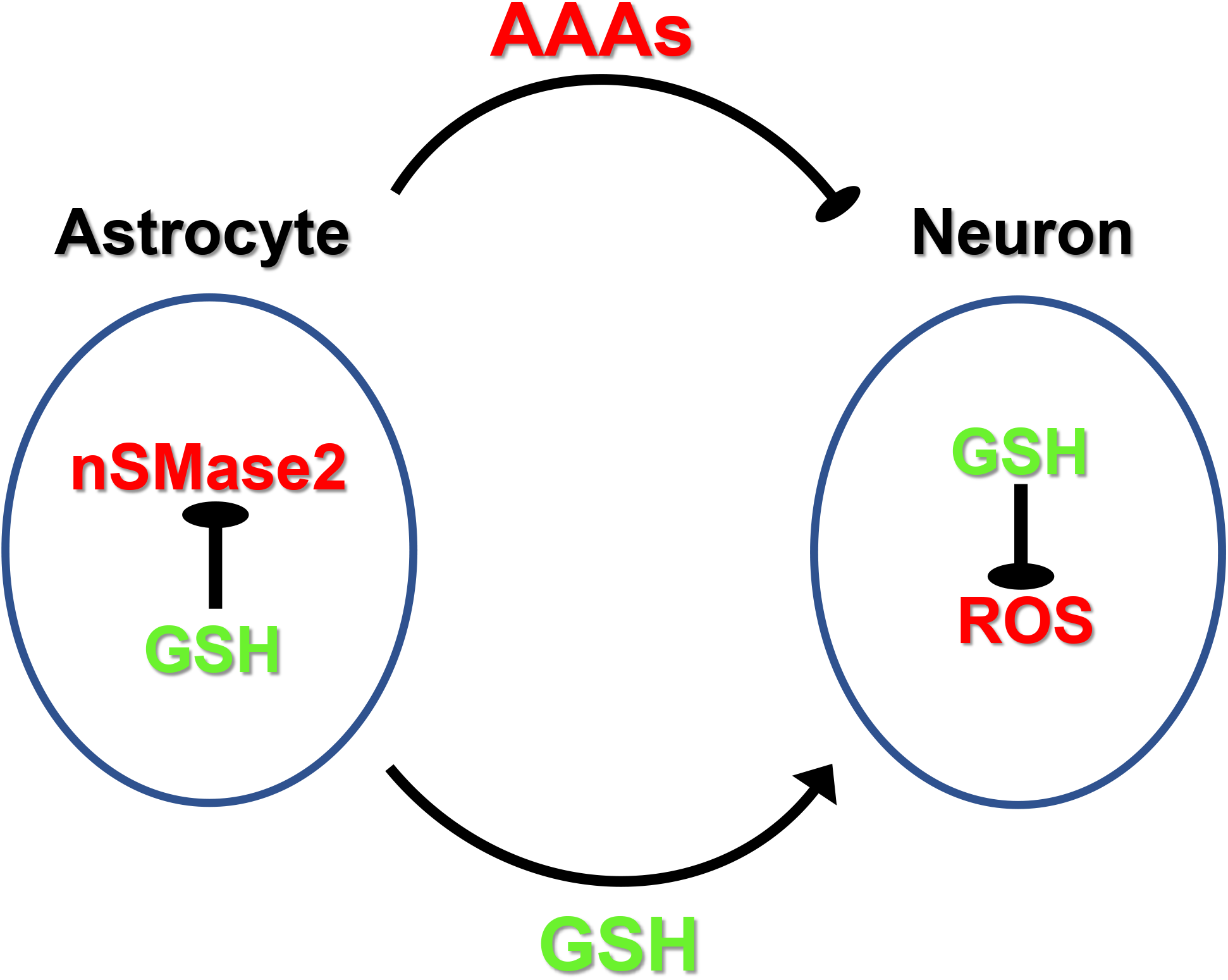
Hypothetical model for regulation of neuronal function by nSMase2 in astrocytes and aging-associated astrosomes (AAAs) Oxidative stress leads to the generation of reactive oxygen species (ROS) in astrocytes and in neurons. In astrocytes, GSH is secreted to provide GSH precursors to neurons. In neurons, GSH resynthesized from astrocytes-derived precursors neutralizes ROS to maintain neuronal function. Increased oxidative stress during aging leads to reduction of GSH levels in astrocytes and activation of nSMase2, which induces formation and secretion of aging-associated astrosomes (AAAs) that suppress neuronal activity. In nSMase2-deficient (*fro/fro*) mice, lack of ceramide elevation prevents formation of AAAs, and in turn, preserves neuronal function.

## Acknowledgements

This work was supported by NIH grants R01NS095215, R01AG034389, and R01AG064234, and the VA grant I01BX003643. The lipidomics analysis of this study was supported in part by the Lipidomics Shared Resource, Hollings Cancer Center, Medical University of South Carolina (P30 CA138313 and P30 GM103339). We thank the Department of Physiology (Chair Dr. Alan Daugherty) at the University of Kentucky, Lexington, KY for institutional support.

